# A Prodrug Strategy to Reposition Atovaquone as a Long-Acting Injectable for Malaria Chemoprotection

**DOI:** 10.1101/2024.02.08.579395

**Authors:** Anil K. Gupta, Anders M. Eliasen, Wil Andahazy, Fei Zhou, Kerstin Henson, Victor Chi, Ashley K. Woods, Sean B. Joseph, Kelli L. Kuhen, John Wisler, Hanu Ramachandruni, James Duffy, Jeremy N. Burrows, Elizabeth Vadas, Andrew Slade, Peter G. Schultz, Case W. McNamara, Arnab K. Chatterjee

## Abstract

Recent malaria drug discovery approaches have been extensively focused on the development of oral, smallmolecule inhibitors for disease treatment whereas parenteral routes of administration have been avoided due to limitations in deploying a shelf-stable injectable even though it could be dosed less frequently. However, an updated target candidate profile from Medicines for Malaria Venture (MMV) and stakeholders have advocated for long-acting injectable chemopreventive agents as an important interventive tool to improve malaria prevention. Here, we present strategies for the development of a long-acting, intramuscular, injectable atovaquone prophylactic therapy. We have generated three prodrug approaches that are contrasted by their differential physiochemical properties and pharmacokinetic profiles: mCBK068, a docosahexaenoic acid ester of atovaquone formulated in sesame oil, mCKX352, a heptanoic acid ester of atovaquone formulated as a solution in sesame oil, and mCBE161, an acetic acid ester of atovaquone formulated as an aqueous suspension. As a result, from a single 20 mg/kg intramuscular injection, mCKX352 and mCBE161 maintain blood plasma exposure of atovaquone above the minimal efficacious concentration for >70 days and >30 days, respectively, in cynomolgus monkeys. The differences in plasma exposure are reflective of the prodrug strategy, which imparts altered chemical properties that ultimately influence aqueous solubility and depot release kinetics. On the strength of the pharmacokinetic and safety profiles, mCBE161 is being advanced as a first-in-class clinical candidate for first-in-human trials.

## INTRODUCTION

Malaria—driven by mosquito transmission of the parasite *Plasmodium spp*.*—*continues to be a significant source of morbidity (241 million annual cases) and mortality (627,000 deaths) within resource-limited countries in Southeast Asia and Sub-Saharan Africa^1^. Since 2015, no significant progress has been made in reducing global malaria cases, particularly in the Democratic Republic of the Congo and northern Nigeria ^2^. Before then, an approximately 40% decrease in malaria between 2000 and 2015 was observed as a result of a World Health Organization (WHO) malaria control program which involved the use of insecticide-treated bed nets (ITNs) and indoor residual spraying (IRS) for vector control, as well as prompt diagnosis and treatment of the infected individuals ^3^. Also, the recommended preventive therapies such as Intermittent Preventive Treatment in pregnancy (IPTp) and Intermittent Preventive Treatment in infancy (IPTi) that rely on sulfadoxine-pyrimethamine (SP) are compromised by drug-resistance parasites, which presents a daunting task for these historical prophylaxis methods to remain effective^4, 5^. However, seasonal malaria chemoprevention (SMC) campaigns targeting children <5 years old in the Sahel region has proven more effective. SMC involves the addition of amodiaquine to sulfadoxine-pyrimethamine (SPAQ) and this regimen is administered monthly throughout the rainy season. Remarkably, this approach has led to a massive reduction in malaria cases by 80% among children^6, 7^. Yet, concerns about SMC include rebound of the disease due to loss of immunity at older ages, unwanted interruptions in campaigns, and development of drug resistance. To address resistance, a new drug combination dihydroartemisinin-piperaquine has been employed to prevent malaria in high-risk groups^8^.

The need for continued development of new therapies to prevent, treat, and control malaria remains paramount to ensure the long-term goals to eradicate malaria by 2030, as outlined in the United Nations (UN) Sustainable Development Goals (SDG). While significant efforts have been made in the development of a biological malaria vaccine (such as RTS, S), the inconsistent, low, and quickly waning efficacy highlights significant challenges^9^. Several options are available for prophylaxis including drugs such as mefloquine (weekly dose), doxycycline (daily dose), atovaquone (ATO)-proguanil (Malarone®; daily dose), and tafenoquine (Arakoda®; weekly dose; recently approved by US FDA), which have been administered to travelers, children, and pregnant women (tafenoquine is not recommended to pregnant women)^10^. However, these options are not ideal due to adverse events, cost, access, resistance, and frequent daily dosing of these drugs. There remains an urgent need for the development of long-acting, oral drugs with once-monthly or less frequent dosing or with a parenteral administration to achieve prophylaxis for at least three months from a single dose.

A wide variety of antimalarial drugs are currently in preclinical and clinical development through MMV and other groups^11-13^: almost all of which are focused on oral administration, with the exception of parenteral administration for severe infections. Subsequently, the question arises whether a known or new anti-malarial compound can be administered via intramuscular (*i*.*m*.) or subcutaneous (*s*.*c*.) routes for long-acting prophylactic treatments of malaria^14^. In the past, there have been considerable efforts to develop injectable depot anti-malarials, but they had limited success. Cycloguanil pamoate along with 4-4′-diacetylaminosulphone (DADDS) was tested in rhesus monkeys using IM injections; however, activity was marred by resistance to pyrimethamine^15-17^. The drug pyronaridine which has been dosed IM once daily for three days (total 480 mg) was shown to be effective for the treatment of chloroquine-resistant *P. falciparum*, and there were no cases of recrudescence observed in all 10 patients even after 30 days^18^. In the past, MMV and Janssen Pharmaceuticals have worked on the development of P218 as a potential long-acting injectable (LAI) chemoprotective medicine^19^. Collectively, the lessons learned from these efforts highlight the need for smaller injection volumes with reduced viscosity (to reduce pain), longer duration coverage from a single administration, and good stability under Zone IV climate conditions.

The concept of an LAI depot is an established approach, as it has been used for the sustained delivery of antipsychotic medicines and hormone therapy^20, 21^. Anti-infectives have also been administered traditionally as injectables of IM penicillin benzathine salts for bacterial infections but with relatively short duration of exposure (14 days with a 1.2 mL injection that can lead to significant injection pain)^22, 23^. Moreover, this approach has more recently been adopted by the HIV field to deliver long-acting formulations of rilpivirine and cabotegravir for pre-exposure prophylactic use (PrEP)^24, 25^. Both of these HIV inhibitor formulations rely upon aqueous suspensions of solid compound nanoparticles that have the advantages of low viscosity and particle size-controlled diffusion rate. However, oil-based solutions have been used for schizophrenia and hormone injection-therapies^20^. Both approaches, aqueous suspensions^26, 27^ and oil-based solutions^28^, are well-accepted, and the physiochemical properties of the active pharmaceutical ingredient (API) generally dictate the formulation approach Alternatively, the API may be re-designed into a prodrug to impart more favorable formulation properties and intrinsic clearance^29^. In general, injection volumes of known LAIs vary from 1-3 mL per injection, and the frequency of injections range from every two weeks to up to 6-month intervals. The injection volume and duration of treatment are principally dependent on the efficacious exposure level that must be achieved, the rate of compound clearance from patients, the dissolution rate of the compound from the depot, and the maximum achievable compound loading in the injection depot. Whereas the compound clearance and the minimal efficacious exposure are intrinsic to the compound, prodrug modulation may be used to manipulate compound stability, solubility and dissolution rate from the depot without changing fundamental biological and pharmacological properties of the parent molecule. Importantly, control of the dissolution rate from the depot can have a profound influence on the overall pharmacokinetic profile, generating a ‘flattened’ profile that restricts the maximal concentration (C_max_) and potentially provides an improved therapeutic safety window. While prodrugs could provide desired profiles needed for optimal formulations and thus required plasma levels for extended protection, the residual circulating prodrugs could cause adverse events and at times, prodrugs might be too stable to get hydrolyzed to provide active drug.

In addition to the modification of API and its prodrug, a third approach could involve slow-release drug delivery systems such as implants. More recently, the Walter Reed Army Institute of Research (WRAIR) has reported development of a long-term malarial chemoprophylactic drug releasing implant and shown the proof-of-concept using piperaquine and ethylene-vinyl acetate (EVA) copolymer^30,31^. In addition to insecticide-treated bed nets for vector control, the use of endectocides has also been considered^32^. Ivermectin, a drug that disrupts malaria transmission by killing mosquitoes, has been formulated as an oral drug capsule that unfurls into a star shape with extended stomach residence time and releases its contents for up to 2 weeks in a pig model^33^. While the former approach, with few exceptions, needs surgery to place and remove the implant, and is likely to be expensive, the latter’s potential is still being explored. New compounds have been identified recently as vector control approaches to prevent malaria ^34^.

MMV recently highlighted the need to extend the chemotherapy armamentarium against malaria to achieve the long-term malaria elimination goals. This was accompanied by changes to the target candidate profiles (TCPs) and target product profiles (TPPs)^35^. In brief, TCP-4 encompasses compounds that minimally possess liver schizonticide activity and are capable of preventing symptomatic malaria. Moreover, it is now acceptable for such compounds with asexual blood stage activity, with or without liver stage potency, to be administered *via* a parenteral route. Either subcutaneous or IM injections are acceptable if they result in long-acting efficacious exposure for ≥3 months; require only a single, small injection volume of <2 mL for adults and <0.5 mL for infants; and can be administered through a 25gauge needle. This approach aligns well for use in chemoprevention/ prophylaxis because a single-dose injection may provide continuous antimalarial prophylaxis for >>1 month. Due to widespread poverty and limited infrastructure in endemic Africa, the cost per injection should be low. The recent revision of the MMV TPPs and TCPs to include parenterally administered chemopreventative/ prophylactic agents underscores a paradigm shift to consider alternative strategies that can further contribute to the control of malaria to reach the larger goals for malaria elimination and eradication.

To develop an LAI for malaria, a suitable known antimalarial drug needs to be selected. In this regard, ATO is a favorable candidate of choice. This compound is a safe and well-tolerated medicine approved for malaria treatment and prophylaxis in combination with proguanil (commercially known as Malarone®); chronic dosing for more than six months has been reported to have no significant adverse effects^36^. In addition to the known safety profile, ATO has exquisite activity against the asymptomatic parasite liver stages and clinical prophylaxis is reportedly achieved with blood plasma concentrations of 100 ng/mL (272 nM) based on a human challenge model^37^. Another report in 2005 suggested concentrations lower than 272 nM could be sufficient for full protection when an ATO-proguanil mixture was administered in humans^38^. A recent study reported 200 ng/mL as the minimal efficacious plasma level for prophylaxis in the *Plasmodium berghei* mouse model.^39^ Furthermore, the relatively long biological half-life (∼2 days) is reflective of low drug clearance and is an essential intrinsic property for the successful development of an LAI. To this end, a recent proof-of-concept study by Bakshi *et al*. described the preparation of ATO nanoparticles in an aqueous suspension capable of maintaining chemopreventative exposure for up to 3 weeks with a single IM injection of a 200 mg/kg dose^39^. They reported that the protection was obtained at plasma concentrations >200 ng/mL (545 nM) of ATO in a malaria mouse model. Interestingly, there has not been any report to date pertaining to the development of ATO prodrugs as LAIs for wider coverage. To this front, another preclinical candidate, ELQ-300 and/or its prodrugs are currently under development for chemoprophylaxis ^40^.

Herein, we describe one-step synthesis of ATO prodrug strategies to (i) preferentially favor oil solution and/or aqueous suspension formulations and (ii) reduce the absorption rate constant of ATO from the depot to extend prophylactic blood plasma concentration. The latter point is critical to induce ‘flip-flop’ kinetics, a state in which absorption rates are slower than the elimination rates. Therefore, the time to steady-state is a function of the absorption rate, and the concentration at steady-state is a function of the elimination rate. Importantly, the hydroxyl group in the quinone ring of ATO offers an opportunistic modification point to attach hydrolyzable prodrug handles to alter the intrinsic physiochemical properties amenable into either an aqueous or oil-based depot formulation. Based on previous human challenge studies, we have used 272 nM as the required minimum efficacious concentration (C_min_) throughout this report (**Figure 1**).

**Figure 1.**
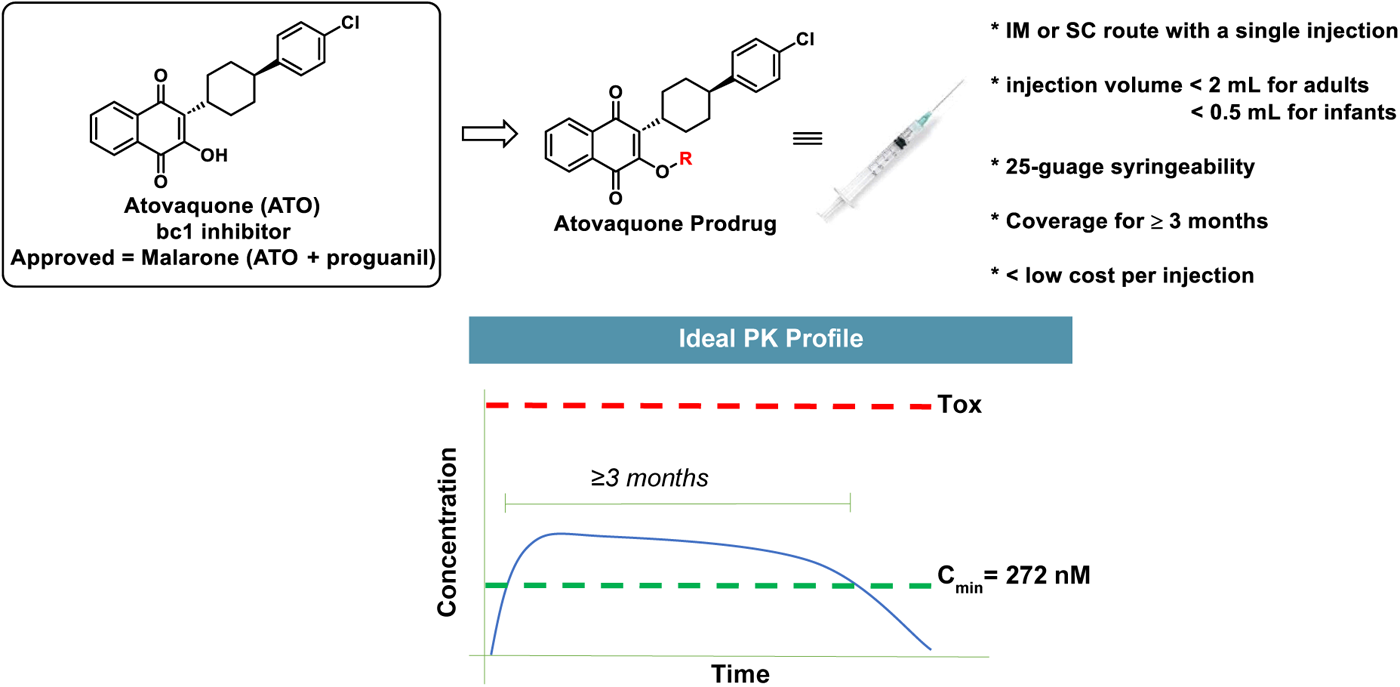
A) Atovaquone (ATO) prodrug strategy. B) Desired PK profile for an ATO prodrug.

## RESULTS

### Limiting Intrinsic Properties of ATO for IM Depot

ATO possesses many requisite properties for an LAI candidate, including low intrinsic clearance, comprehensive and well understood human tolerability profile from ample historic data and exposure data (which suggest <300 nM blood plasma concentrations are prophylactic^37, 38^), and approval by a stringent regulatory agency. While previous efforts have been made to improve oral delivery of ATO^41^, there has not been the same rigorous effort to design an LAI ATO other than the lone example reported recently by Bakshi and co-workers^39^. Previous oral ATO administration determined that protection was provided for 24 days based on the minimum efficacious plasma concertation (i.e., 272 nM; the prophylactic concentration in humans).^37^ Therefore, as a control experiment, a high dose of 75 mg/kg of ATO was orally administered in rats as a solution in 10% M-pyrol/10% ethanol:cremophor 50:50/, 80% D5W (**Fig. 2**). Subsequently, the rat pharmacokinetic (PK) profile of native ATO was evaluated at 20 mg/kg as a 200 mg/mL aqueous suspension in 1% (w/v) F108 and 0.2% (w/v) sodium laureth sulfate (SLS) administered via a single IM injection. Impressively, the IM depot of ATO maintained prophylactic plasma exposure at or above 272 nM for approximately 3 weeks (**Fig. 2**). These results show that ATO can be dosed IM at almost a quarter of the oral (PO) dose for similar exposure (20 vs 75 mg/kg, respectively).

**Figure 2.**
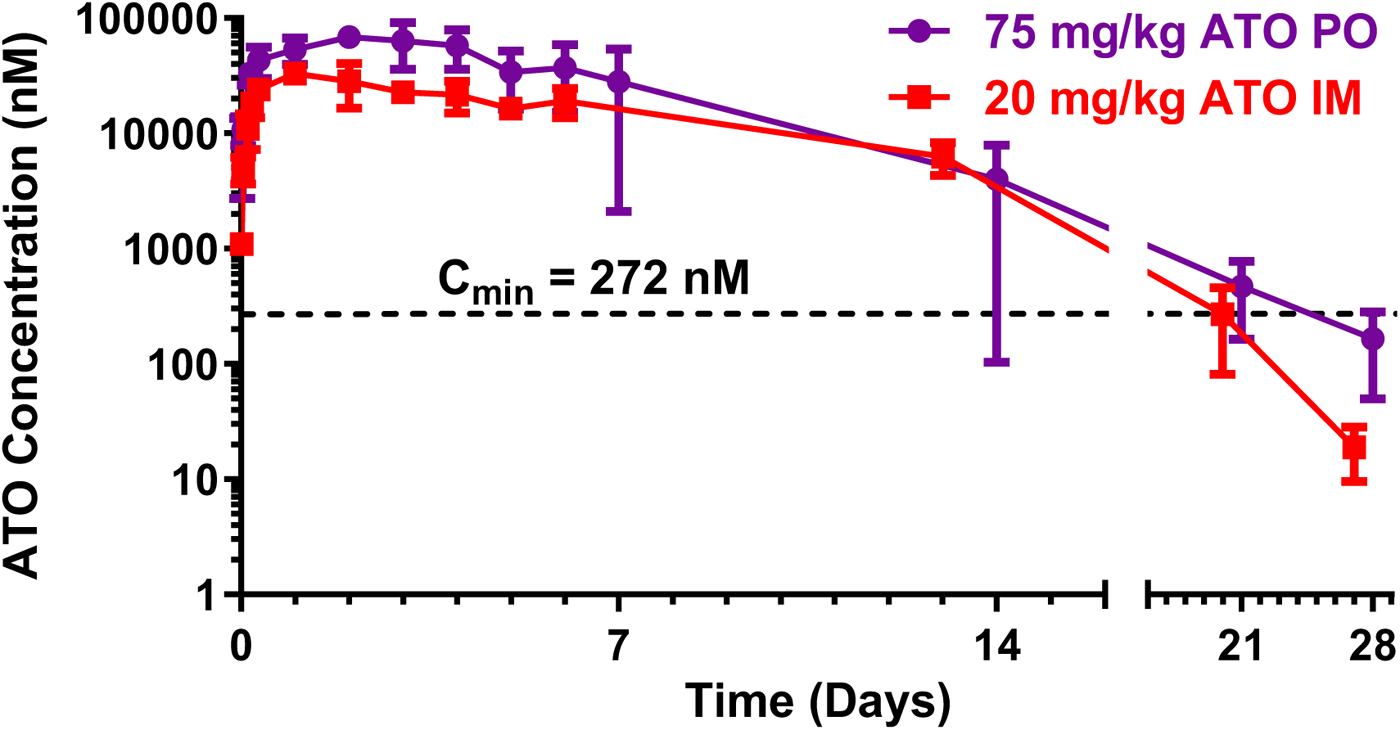
IM ATO provides minimum efficacious plasma exposure at nearly a quarter of the oral dose of ATO. ATO was administered PO or IM in aqueous formulations, and plasma was collected at various timepoints over 28 days. The mean (±SD, n=3) plasma concentration of ATO measured and graphed as a function of time. C_min_=minimal efficacious concentration (272 nM).

However, the current PK profile of IM ATO is suboptimal for a long-acting parenteral injection, highlighted by the “burst” of ATO plasma exposure observed within the first week following administration and the relatively short exposure time of 24 days (**Fig. 2**). Notably, the C_max_ is observed in the first 48 hours after injection, followed by a more rapid elimination until the true equilibrium between depot dissolution and ATO clearance is achieved. It must be noted that an initial burst (higher release rate) may be beneficial for cure of an existing infection as long as an overall duration of the LAI is not significantly affected without any safety concerns. Optimized burst effect offers the advantage of flattening the PK profile from the depot, leading to reduced blood plasma concentration throughout the duration of depot release, and the potential for a significant extension of prophylactic protection.

### Strategy and Prodrug Design for IM ATO Depot

The analysis of the IM ATO depot revealed the need to alter the physiochemical properties of ATO to achieve a more desirable rate of dissolution from the IM depot and promote a flattened PK profile. As a general strategy and, if available, an alcohol or vinylogous amide functional group in the parent compound is modified to an aliphatic ester. The hydroxy in the quinone ring of ATO is a convenient site for prodrug modification with an ester-based linkage. A review of previous ATO prodrugs revealed successful targeting of this same hydroxyl site improved the oral absorption of ATO^42-45^, yet this approach had not been applied to ATO prodrugs for LAIs.

In the case of ATO, a high-yield, one-step synthesis was used for all prodrugs (**Fig. S1**). Among all prodrugs, final optimized compounds mCBE161, mCKX352, and mCBK068 derived from acetic acid, heptanoic acid, and docosahexaenoic acid, respectively, were generated using a large span in the number of carbon atoms in the aliphatic group (**Table 1**). The properties of the aliphatic group control the solubility and dissolution of the prodrug from the depot site into the bloodstream, whereas the prodrug encounters ubiquitous hydrolases in the blood plasma that rapidly cleave the ester bond and revert the antimalarial prodrug into its parent compound. Aliphatic groups classified as generally recognized as safe (GRAS) were prioritized as preferred ester components to avoid the unnecessary introduction of potentially toxic organic acids following hydrolysis. Docosahexaenoic acid (DHA; hydrolysis product of mCBK068) and heptanoic acid (hydrolysis product of mCKX352) are exemplars of this strategy. The DHA and heptanoyl chains impart a high calculated log partition (clogP) value to their respective prodrugs mCBK068 and mCKX352. The resulting high lipophilicity makes these molecules suitable candidates for oil solution formation. mCBK068 is derived from DHA which is the longest omega-3 fatty acid and possesses the highest degree of unsaturation in this family and is GRAS. The DHA-derived ATO ester mCBK068 was an oil, whereas the heptanoate ester mCBK352 was found to be a crystalline solid with a low 76.1°C melting point. mCBE161, similar to ATO, was also found to be crystalline solid with a relatively higher melting point than other prodrugs.

**Table 1.**
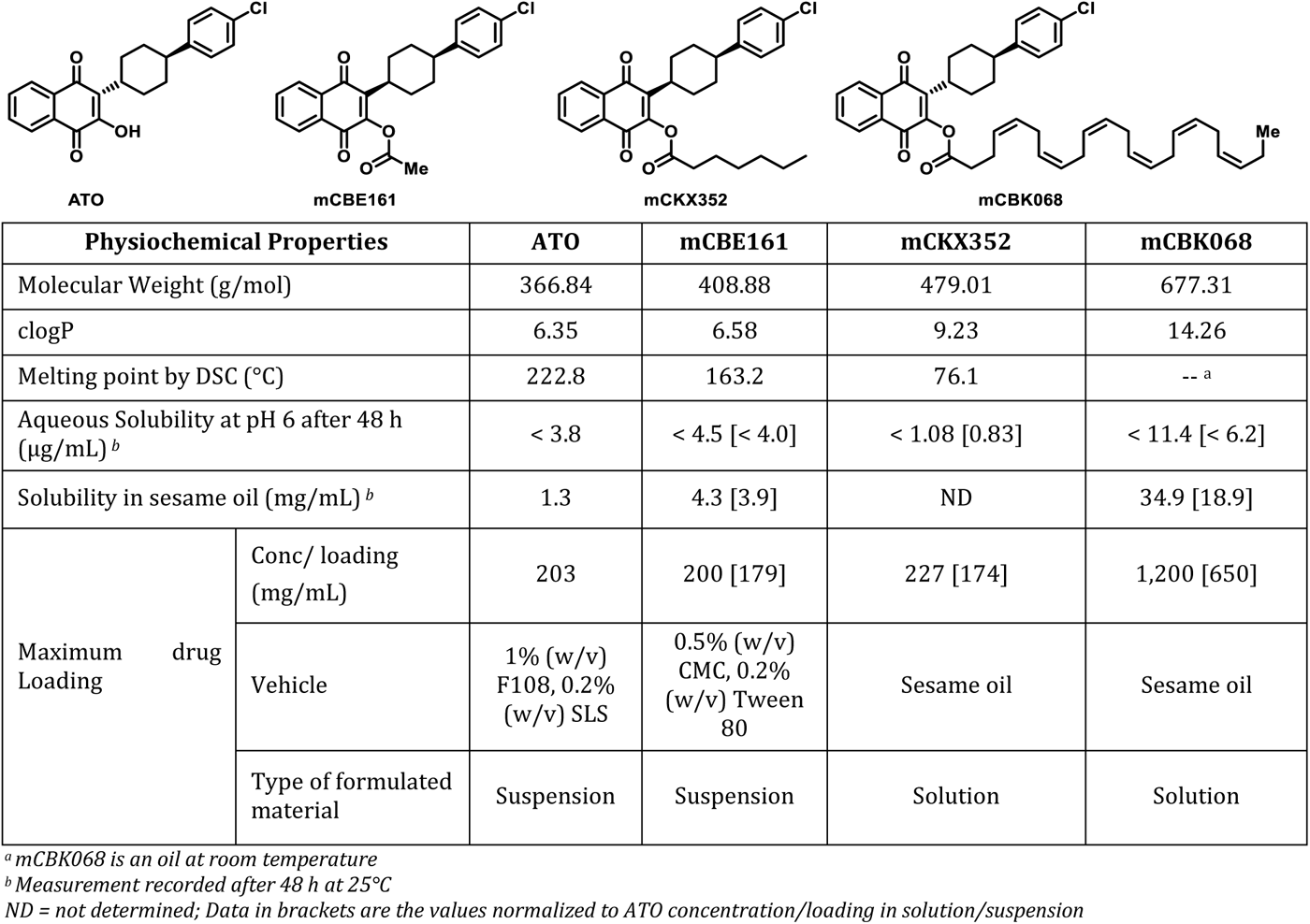
Summary of physical properties of ATO and prodrug lead candidates.

All prodrugs, in addition to ATO, have very low aqueous solubility. Each remain neutral across the physiologically relevant pH range and as a result, the solubility is essentially independent of pH. While the physiochemical properties such as high clogP/LogD and high solubility in various clinically acceptable oils are consistent with an oil depot for mCKX352 and mCBK068, properties such as high melting point and low aqueous/oil solubility, as observed for mCBE161 and ATO, are suitable for IM administration either as an oil or aqueous suspension depot (see **Table S1** for details).

### Rat PK Studies of IM Depots of ATO Prodrugs

Similar to ATO, mCBE161, an acetic acid ester prodrug of ATO, was IM administered as an aqueous suspension (0.1% (w/v) F108 and 0.2% (w/v) SLS) of 20 mg/kg (∼18 mg/kg ATO after accounting for the prodrug mass) at a drug loading of 240 mg/mL (215 mg/mL ATO). Interestingly, mCBE161 provided sustained mean blood plasma exposure of ATO that was above the minimal efficacious concentration of 272 nM for approximately 69 days (**Fig. 3**). Examination of the C_max_ also showed a desired reduction for mCBE161 (C_max_ = 11,149 nM), which is approximately one third of the concentration for the aqueous suspension of ATO (35,532 nM) (**Fig. 3**).

**Figure 3.**
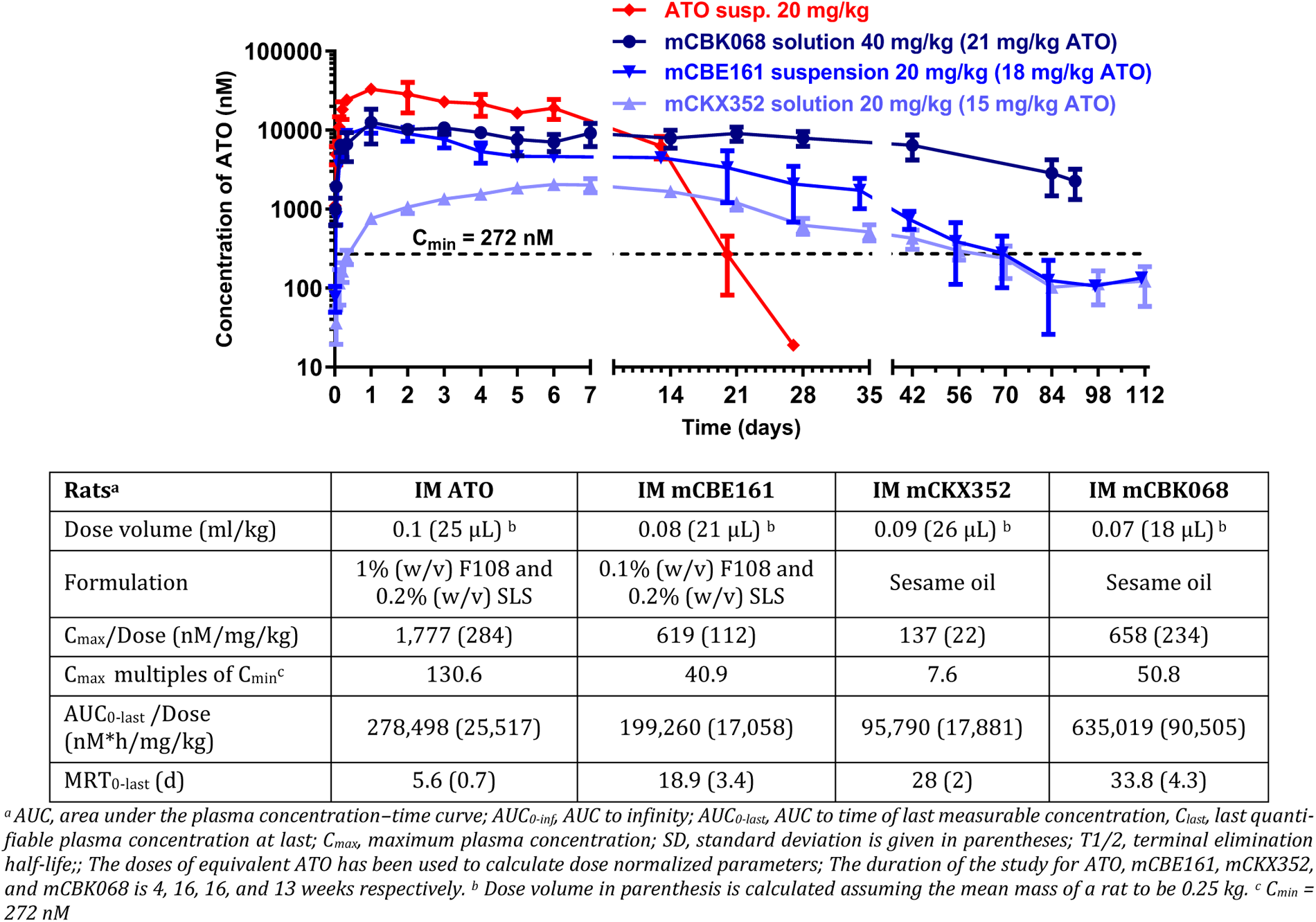
IM ATO prodrugs have longer minimum efficacious exposure with less drug than IM ATO. (**Graph**) 4-month rat PK profile for IM administered ATO and ATO prodrugs. The mean (±SD, n=3) plasma concentration of ATO as a function of time following IM injection of ATO (aqueous suspension), mCBE161 (aqueous suspension), mCKX352 (oil solution), and mCBK068 (oil solution) in rats. (**Table**) Summary of key rat PK measurements for ATO, mCBE161, mCKX352, and mCBK068 (see **Table S2** for details). C_min_=minimal efficacious concentration (272 nM).

Based on the relatively high oil solubility, mCBK068 and mCKX352 were chosen to advance to *in vivo* PK profiling as an oil solution depot in rats.

While mCKX352 was dosed at 20 mg/kg (∼15 mg/kg ATO) with a concentration of 227 mg/mL (174 mg/mL ATO) and dose volume of 0.09 mL/kg, the IM dose of mCBK068 was 40 mg/kg (∼21 mg/kg ATO) at a concentration of 545 mg/mL (295 mg/mL ATO) with dose volume of 0.07 mL/kg. While mCKX352 provided sustained plasma exposure of ATO above the minimal efficacious concentration of 272 nM for up to 56 days, the coverage exceeded 4 months in the case mCBK068 with C_last_ = 2,256 nM which is 8-fold higher than the minimal efficacious exposure.

Consistent with the mCBE161 depot, the C_max_ of the oil solution IM depots of mCKX352 (2,058 nM) and mCBK068 (13,812 nM) are significantly lower than that for the aqueous suspension of ATO (35,532 nM). Correspondingly, compared to the aqueous suspension depot, the plasma concentration profile remains significantly ‘flattened’ and sustained for rats administered the mCKX352 and mCBK068 oil-based solution depot. The C_max_/C_min_ window has been observed to be as low as 7.6 proving the advantage of prodrugs to have an optimized burst of ATO (**Fig. 3**). Henceforth, The PK profile highlights a key differentiating feature of the ATO prodrug strategy. Here, the prodrugs mCKX352 and mCBK068 appear to provide a slower rate of dissolution from the depot and into the bloodstream. Expectedly, this is contrasted by a considerable high burst of compound dose dumping of ATO when the parent compound is prepared as a simple aqueous suspension (**Fig. 2**). Finally, ATO suspension reached the maximum plasma level within 1.3 days whereas mCKX352 and mCBK068 took approximately 7 and 11 days, respectively. These oil-based solution rat PK profiles showed a 14-fold extended half-life of mCKX352 in comparison to the aqueous suspension of ATO. A similar observation was made for mCBK068. Clearance in the rat was 0.12 mL/min/kg for mCBK068, the lowest clearance observed for any prodrug in the ATO series. Importantly, the prodrug levels of mCKX352 were monitored and found to be less than 50 nM across all time points.

### Formulation Development of Drug Substances to Generate Drug Products

The ideal threshold of an adult dose is <100 mg (or 2 mg/kg) for a preclinical oral antimalarial^46^. To keep injection volumes low (such as <1 mL), the minimum acceptable concentration/loading of ATO in any formulation for an IM injectable drug is ∼100 mg/mL.

mCBE161 was initially formulated as a simple aqueous suspension generated from F108 and SLS for evaluation in rat PK. However, since SLS is a potential skin irritant, an alternate aqueous suspension formulation involving methylcellulose derivatives and Tween 80 was explored. This led to the generation of the well-behaved aqueous suspension in 0.5% (w/v) carboxymethyl cellulose (CMC) and 0.2% (w/v) Tween 80. Similar to the F108/SLS-based suspension, the maximum drug loading of mCBE161 is 200 mg/ml (equivalent to ∼179 mg/mL ATO), which permits a smaller injection volume (**Table 2**).

**Table 2.** Formulation development of ATO prodrug lead compounds.

| Compounds | Maximum loading or concentration of Prodrug (mg/ml) | Maximum loading or concentration of ATO (mg/ml) | Formulation | Type of formulation |
| --- | --- | --- | --- | --- |
| ATO | Not Applicable | 203 | 1% (w/v) F108, 0.2% w/v SLS | suspension |
| mCBE161 | 246 | 221 | 0.1% (w/v) F108, 0.2% w/v SLS | suspension |
|  | 231 | 207 | 0.5% (w/v) MC, 0.2% (w/v) Tween 80 | suspension |
|  | 200 | 179 | 0.5% (w/v) CMC-Na, 0.2% (w/v) Tween 80 | suspension |
| mCKX352 | 227 | 174 | sesame oil | solution |
| mCBK068 | 1,200 | 650 | sesame oil | solution |

CMC=carboxymethylcellulose; MC=methylcellulose; SLS=sodium laureth sulfate.

Solubility studies of mCKX352 were conducted in various clinically acceptable oils including fractionated coconut oil, soybean oil, olive oil, and castor oil at 25ºC for up to 7 days. Solubility of mCKX352 in all four oils remained stable for 7 days without visible precipitation or other physical changes (see Supplementary Materials for details)

The presence of the longest omega-3 fatty acid chain in mCBK068 resulted in maximum concentration of ATO as high as 650 mg/mL in sesame oil (**Table 2**). It should be noted that the DHA ATO ester is an oil that is syringeable in a 25-gauge needle and remains miscible in sesame oil.

### Stability Studies of Drug Substances and Drug Products

Regarding chemical and pH stability, all prodrugs were stable under ambient temperature and were subjected to stability studies in pH 2/4/6/8 buffers (50:50, MeCN:H_2_O) for up to 7 days (**Table 3**). All compounds were >95% pure by HPLC-UV and HPLC-MS analyses. For ATO, no obvious degradation was observed in any of the pH buffers for 7 days. For mCBE161, significant degradation (>10 area% decrease) was observed in pH 8 buffer for 2 hours, and slight degradation (∼5.9 area% decrease) occurred in pH 6 buffer for 7 days, and a slight increase in purity was observed in pH 2/4 buffers for 7 days possibly due to further degradation of certain impurities. For mCKX352, while significant degradation (∼60 area% decrease) was observed in pH 8 buffer for 1day, slight degradation (∼2.1 area% decrease) occurred in pH 6 buffer for 7 days. Also, a slight increase in purity was observed in pH 2/4 buffers for 7 days possibly due to further degradation of certain impurities. For mCBK068, significant degradation (>30 area% decrease) was observed in pH 8 buffer for 1 day, and no obvious degradation occurred in pH 4/6 buffers for 7 days. Finally, a slight increase in purity was observed in pH 2 buffer for 7 days possibly due to further degradation of certain impurities.

**Table 3.** Accelerated stability evaluation of ATO prodrug lead compound.

| Conditions | mCBE161<br>(suspension in 0.5% (w/v) CMC, 0.2% (w/v) Tween 80) |  |
| --- | --- | --- |
| | Purity (%) after 1 week | Average particle size (μm) |
| Initial <sup>a</sup> | NA | 2.1 |
| 25 °C/60% RH | 99.4 <sup>b</sup> | 2.7 |
| 40 °C/75% RH | 99.3 | 3.3 |
| 60 °C/ambient | 99.2 | 3.0 |

<sup>a</sup> Purity measured directly after preparation
<sup>b</sup> Relative humidity was not controlled
RH = Relative Humidity; NA = Not Available; ND = Not Determined

All mCBE161 formulations were subjected to accelerated stability studies required for clinical development (**Table 2**). All formulations were stable, and particularly, the suspension with 0.5% (w/v) CMC-Na, 0.2% (w/v) Tween 80 was physically and chemically stable under three conditions (RT, 40 °C/75% RH, 60 °C) for 7 days, and mean particle size is ∼2-3 μm (**Table 3**). Importantly, the resulting aqueous suspension maintained a sufficiently low viscosity for use of a 25-gauge syringe, which is an ideal requirement for IM delivery of these prodrugs. In parallel, mCBE161 stability studies were also performed in sesame oil and Miglyol 812, which exhibited slight degradation (<2.7 area% decrease) under three conditions (25°C/60% RH, 40°C/75% RH and 40°C/ambient) for up to 4 weeks. This was done for a possible oil-based suspension development if the aqueous-based suspension formulation approach were to encounter unforeseen limitations in subsequent development.

### PK Studies of Cynomolgus Monkey IM Depots

After rat PK analysis, the focus was shifted to non-rodent species. Based on the previous studies reported for ATO, dog was selected as the preferred species and PK was performed using male beagle dogs. The dog PK of mCBK068 exhibited an unexpectedly different PK profile than that observed in the rat. This was characterized by a very delayed time to peak drug concentration (T_max_) value ranging from 18 to 27 days in all routes and doses; and henceforth, it was determined that the dog muscular structure may be different fromthat of rat. Therefore, mCBK068 was advanced for cynomolgus monkey (cyno) PK, which ultimately correlated very well to the observed rat PK. For the IM dose of 19 mg/kg (∼10 mg/kg ATO) at a concentration of 190 mg/mL (103 mg/mL ATO), the formulation was 100% sesame oil and administered IM at the left/right muscular flank (biceps femoris) of monkey (**Fig. 4**). mCBK068 provided sustained plasma exposure of ATO above the minimal efficacious concentrations of 272 nM for up to 41 days. The mean residence time was approximately 1.5 months with C_max_ reached on day 5. At all points, prodrug mCBK068 was found to be below the quantitation limit (BQL). Subsequently, cyno PK was initiated for mCKX352 and mCBE161 using male cynomolgus monkeys with three monkeys per group. While mCKX352 was dosed with 13 mg/kg (∼10 mg/kg ATO) at a concentration of 100 mg/mL (77 mg/mL ATO) in sesame oil, the IM dose of mCBE161 was 11 mg/kg (∼10 mg/kg ATO) at a loading of 100 mg/mL (90 mg/mL ATO) as an aqueous suspension using 0.5% (w/v) CMC - Na /0.2% (w/v) Tween 80. It is important to mention that the formulation for mCBE161 was modified to remove SLS because of its potential as a possible skin irritant. For mCKX352, a flattened PK curve with no dose dumping provided plasma levels higher than both C_min_ values for >70 days. This result is different from the previous oil-based solution of mCBK068 which provided coverage for 41 days for C_min_ = 272 nM. mCBE161 also afforded excellent exposure for >30 days which was higher than C_min_. Unlike mCBK068, both mCBE161 and mCKX352 have significantly delayed T_max_ values, which were reached after two weeks. The oil-based solution rat PK profile of mCKX352 resulted in an extended mean residence time and lower C_max_ than that observed for the aqueous suspension of mCBE161. The C_max_/C_min_ window was lowered to ∼3–4 for oil solutions, which is almost 3fold better than the aqueous suspension formulation (**Fig. 4**). Across all time points, similar to the rat PK results, the prodrug levels of mCKX352 and mCBE161 were found to be less than 15 nM and BQL, respectively. Importantly, in all cases, due to the ability to achieve high prodrug concentration within the formulation, low dose volumes in the range of 290–430 uL were injected. Based on instability of the drug product from mCBK068 and relatively lower coverage in cyno PK, it was decided to move forward with mCKX352 as our oil-based prodrug in subsequent two-week rat toxicology studies.

**Figure 4.**
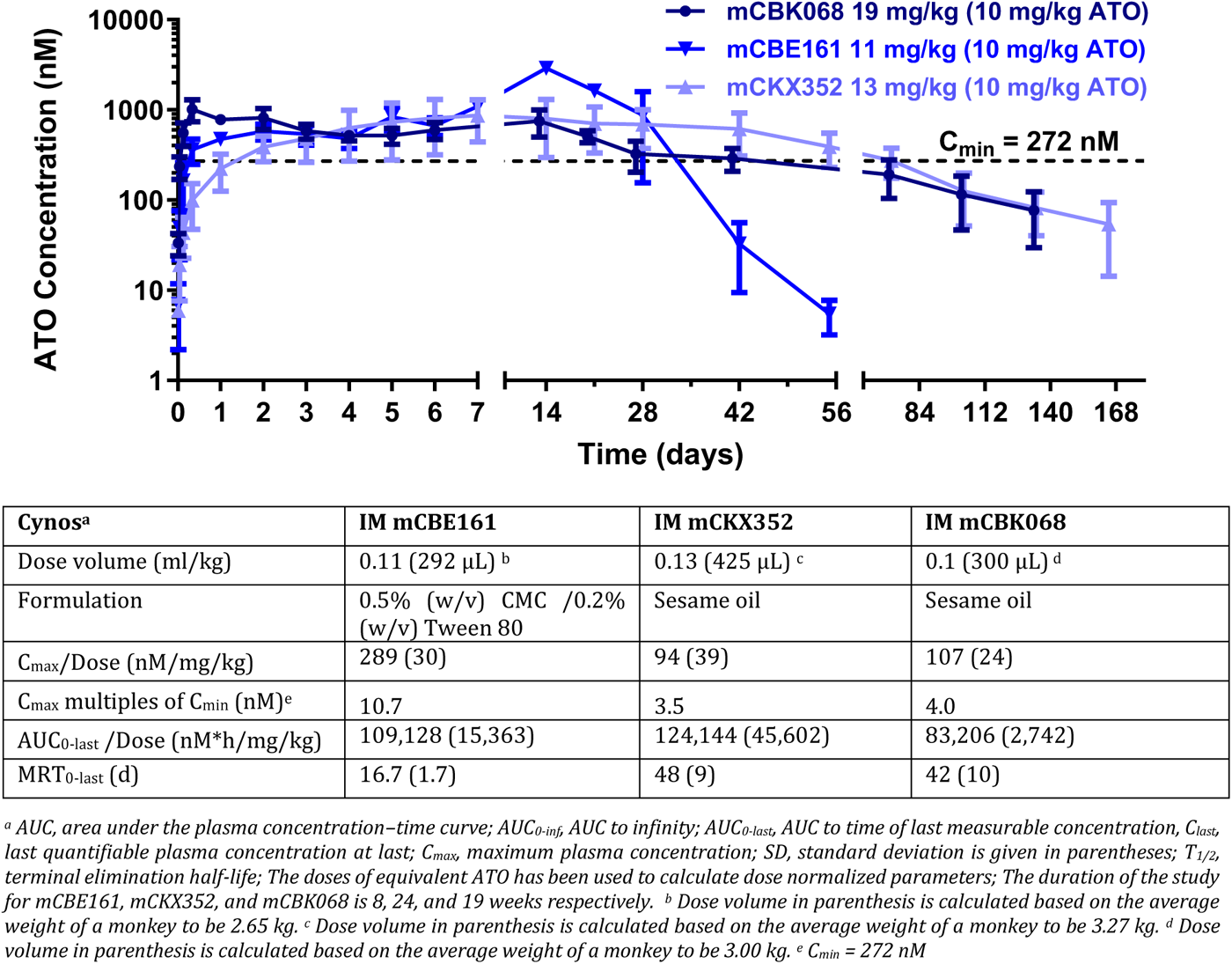
IM-administered ATO prodrugs show wide variations in exposure time in cyno PK studies. (**Graph**) 6-month cyno PK profile for IM-administered ATO prodrugs. The mean (±SD; n=3) plasma concentration of ATO as a function of time following IM injection of mCBE161 (aqueous suspension), mCKX352 (oil solution), and mCBK068 (oil solution) in cynos. (**Table**) Summary of key rat PK measured for mCBE161, mCKX352, and mCBK068 (see **Table S2** for details); C_min_=minimal efficacious dose based on ^37^.

### Rat Toxicology and Toxicokinetic (TK) Studies

Both prodrugs mCBE161 and mCKX352 were evaluated in a 2-week rat toxicity studies via IM administration. In the mCBE161 study, the vehicle for dose formulation preparation was 0.5% CMC & 0.2% Tween 80 in water. Sesame oil was used as the vehicle for dose formulation preparation in the mCKX352 study. Toxicology and TK were assessed in both studies.

In a 2-week non-GLP toxicology study of mCBE161, male and female Sprague Dawley rats (5/sex/group) were dosed via IM injections on days 1 and 14 with vehicle, 20, 80, or 250 mg/kg (18, 72, or 225 mg/kg ATO). TK was evaluated in the vehicle group (3 animals/sex) and in mCBE161 treated groups (6 animals/sex). No gender difference in exposure of mCBE161 were observed. No clinical signs of toxicity (no mortality, body weight, food consumption, etc.) were observed, and all animals survived until the end of the study. Very slight edema and mild swelling at injection sites were found on day 15 at ≥ 20 mg/kg in both genders as well as in female animals in the control group. Minimal-to-moderate inflammation at the injection sites was characterized by presence of mixed inflammatory cells (neutrophils, macrophages, lymphocytes and/or giant cells) with occasional muscle degeneration/necrosis at ≥80 mg/kg/day observed. Based on study results, IM injection did not result in any adverse findings. Therefore, the no observed adverse effect level (NOAEL) of mCBE161 was determined to be >250 mg/kg (225 mg/kg ATO) of mCBE161, which corresponded to day 14 mean C_max_ of 0.57 μg/mL mCBE161 (14.37 μg/mL ATO) and mean AUC value of 110.64 μg*hr/mL (2,751.7 μg*hr/mL ATO).

In another 2-week non-GLP rat toxicity study of mCKX352, male and female Sprague Dawley rats (5 sex/group) were dosed via IM injections on days 1 and 14 with vehicle, 20, 100, or 200 mg/kg mCKX352 (15, 77, or 153 mg/kg ATO). TK was evaluated in the vehicle group (3/sex) and in mCKX352 groups (6/sex). No gender difference in exposure of mCKX352 was observed. No clinical signs of toxicity were found, and all animals survived until the end of the study. Microscopically, mild-to-moderate inflammation was observed at the injection sites characterized by presence of mixed inflammatory cells (neutrophils, macrophages, lymphocytes, and/or giant cells) with occasional muscle degeneration/necrosis at ≥100 mg/kg/day. Skeletal muscle necrosis was observed in the left thigh muscles that received the most recent injection (day 14) but was absent in the right thigh muscles that received the earlier injection on day 1 in the 20 and 100 mg/kg dose groups, which indicate that this finding was reversible. Based on study results, IM injection did not result in any adverse findings. Therefore, the NOAEL was >200 mg/kg of mCKX352 (>153 mg/kg of ATO). At 200 mg/kg, day-14 TK of mCKX352 mean C_max_ was 5.36 μg/mL, and the mean AUC value was 794.29 μg*hr/mL. Subsequently, at 200 mg/kg, day-14 TK of ATO (derived from mCKX352) mean C_max_ was 4.06 μg/mL and the mean AUC values 423.51 μg*hr/mL.

The therapeutic index (TI) for both mCBE161 and mCKX352 based on C_max_ and C_last_ relative to 100 ng/ml (efficacious concentration) is shown in **Table 4**. Based on the C_last_ on day 14, the TI of mCBE161 is 51, 104, and 126 at 20, 80, and 250 mg/kg, respectively. Similarly, the TI of mCKX352 is 4, 9.5, and 16.5 at 20, 100, and 200 mg/kg, respectively. Due to these safety and PK profiles, mCBE161 is being advanced to first-in-human clinical trials.

**Table 4.** Two-week rat IM study therapeutic index (TI) on day 14.

| Prodrug | Dose (mg/kg) | $C_{max}$ (ng/ml) <sup>a</sup> | $C_{last}$ (ng/ml; 240 h) <sup>a</sup> | TI<sup>b</sup> $C_{max}$ | TI<sup>b</sup> $C_{last}$ |
| --- | --- | --- | --- | --- | --- |
| **mCBE161** | 20 | 5,415 | 5,095 | 54.2 | 51 |
|  | 80 | 10,370 | 10,370 | 103.7 | 103.7 |
|  | 250 | 14,370 | 12,625 | 143.7 | 126.3 |
| **mCKX352** | 20 | 925 | 400 | 9.3 | 4 |
|  | 100 | 3,120 | 950 | 31.2 | 9.5 |
|  | 200 | 4,060 | 1,670 | 40.6 | 16.5 |

<sup>a</sup> Average of males and females
<sup>b</sup> Based on $C_{max}$ or $C_{last}$ divided by 100 ng/ml (minimum efficacious concentration)

## DISCUSSION

The multiple goals of LAI malaria chemoprophylaxis include protection of a) children in areas of high seasonal or even perennial transmission, b) migrant populations within Africa traveling into endemic regions; and c) static populations from a resurgent epidemic to “maintain zero” infections. LAI chemoprophylaxis should include easy delivery with low dosing volume to reduce injection site pain, infrequent dosing to improve compliance, and an exceptional local and systemic safety profile. ATO has some of the key attributes that make it a very attractive candidate for a LAI. This includes significant information on systemic human safety and prophylaxis efficacy^36, 47^. One key drawback of ATO compared to ELQ300^40^ and more contemporary cytochrome bc1 inhibitors is the emergence of clinical resistance after short periods of dosing^12, 48, 49^. Therefore, drug combinations such as ATO plus proguanil (Malarone) have been used to address resistance, enabling a first-line oral prophylaxis for many years.

In 1999, data from human malaria challenge studies testing ATO alone suggested a targeted concentration of ≤100 ng/mL (≤272 nM) of ATO is required for complete protection^37,36^. According to subsequent human challenge studies, a protective level^50,49^ of ATO in an ATO-proguanil mixture suggests longer-half-life and overall higher plasma levels (in terms of AUC and C_max_) of ATO led to prophylactic success with C_max_ as low as ∼150 ng/mL (409 nM) of ATO. Due to poor oral bioavailability of ATO, it is challenging to regulate plasma levels unless parenteral administration is employed.

While studies have shown that ATO-resistant *P. falciparum* mutations cannot be transmitted by mosquitoes^51^, the possibility of pre-existing mutations in the parasite population may pose a drug resistance threat for ATO monotherapy. Hence, it will be imperative to identify a combination partner for the continued development of ATO-based LAI strategies. ELQ-300 is an obvious partner candidate, and it has previously been shown to have excellent efficacy in combination when dosed orally in a mouse malaria infection model^52^.

The development path of an injectable antimalarial for chemoprophylaxis is established based on the physiochemical properties, available human safety and efficacy data, and prior approval by a stringent regulatory authority. In a review of the literature for prior strategies to generate an LAI of ATO, only one publication was identified: Bakshi *et al*. demonstrated that an aqueous suspension of ATO nanoparticles could achieve ∼1-month of prophylactic exposure^39^. This approach required a high dose of ATO (200 mg/kg) because of the rapid dissolution of the nanoparticles from the depot. In order to achieve a slow release of ATO from the depot and to generate the desired plasma exposure profile that will support at least three months of chemoprevention from a single dose, a prodrug strategy^53^ has been employed to reduce intrinsic clearance and/or reduce rate of dissolution from the depot site. Alteration of the physiochemical properties of ATO are necessary to reduce the dissolution rate and avoid “dose dumping” from the depot. By instituting a slow release from the depot, it will also favorably optimize the “burst effect,” which we found to be characteristic of IM depot injections. In general, immediately after the injection, a C_max_ is achieved within the first 48 hours, followed by a more rapid elimination until the true equilibrium between prodrug dissolution and ATO clearance is achieved. Minimization of the C_max_ offers two important advantages: 1) flattening of the PK profile from the depot and potential to significantly extend chemoprotective exposure in the blood plasma and 2) a significant increase in the therapeutic window for C_max_-related adverse events.

The hydroxy in the quinone ring of ATO offers an opportunistic modification point to attach hydrolyzable prodrug handles to alter the intrinsic physiochemical properties for depot into either aqueous or oil-based depot. As discussed earlier, there have been efforts to utilize the prodrug strategy to improve the absorption for oral delivery in the case of ATO^42-45^. In addition, new analogs were reported that contained long-chain and short-chain ester prodrugs of ATO^54^. However, these analogs were only tested for in vitro potency and no further characterization of the injectability of these analogs was reported. Altogether, 22 prodrugs of ATO were synthesized and characterized for compound solubility, rat PK plasma exposure, and preliminary formulation stability.

Well-established formulations for LAI include oil solutions, aqueous micro-and nano-suspensions, and polymeric barriers such as in situ-forming gels and microspheres. A list of clinically acceptable solubilizing excipients are water-insoluble lipids such as castor oil, corn oil, cottonseed oil, olive oil, peanut oil, peppermint oil, safflower oil, sesame oil, soybean oil, hydrogenated vegetable oils, hydrogenated soybean oil, and fractionated coconut oil and palm seed oil^55^. Sesame oil has been the oil of choice for LAI for antipsychotics due to its tolerability and optimal viscosity^20^. Henceforth, sesame oil was chosen for all the oil solution PK studies. mCBK068 was the first advanced lead as an oil-based prodrug to emerge from the initial workflow; however, inherent instability of the drug product containing docosahexaenoate chain could not be mitigated. Replacement of DHA with heptanoic acid in mCKX352 helped overcome the limitation of the previous lead candidate mCBK068. Elimination of the unsaturated bonds and introduction of a completely saturated aliphatic chain demonstrated much improved stability under accelerated conditions.

The second most common and well-adapted LAI formulation is aqueous micro- and nano-suspensions. A better understanding of the particle size and dissolution rate aids in optimizing the release of the drug/prodrug, thereby increasing the duration of plasma concentrations of the drug up to the desired level. The major disadvantages of the aqueous suspension are relatively higher cost of goods than oil solutions and complex development, making reproducibility a critical problem. Nonetheless, based on the lower melting point as compared to ATO and low aqueous solubility, mCBE161 was chosen to advance to *in vivo* PK profiling as an aqueous suspension. The final formulation for the cyno PK included Na-CMC, which is a well-accepted excipient approved for IM dosing^20^. The maximum concentration of mCBE161 obtained was 200 mg/ml (∼179 mg/mL ATO) when an aqueous suspension was prepared using 0.5% (w/v) CMC-Na, 0.2% (w/v) Tween 80. Henceforth, the high suspension loading of mCBE161 will ensure ideal injection volumes less than 1 mL.

To avoid an expensive cold-chain requirement in resource-limited areas, the drug product needs to be stable and have >2-5 years of shelf-life according to Zone IV stability standards^56^. As discussed earlier, mCKX352 showed excellent stability under accelerated conditions in clinically accepted fractionated coconut oil. Even in the case of aqueous suspension of mCBE161, no significant increase in particle size leading to Ostwald ripening^57^ was observed under accelerated conditions; Smaller particle size (higher surface area) gives optimal dissolution rate without clogging the needle.

The subsequent head-to-head profiling of the aqueous suspension for ATO and the ATO prodrug mCBE161 demonstrated superior dissolution kinetics of the latter from the depot. This resulted in an LAI profile with efficacious plasma concentrations for more than 2 months with a 20 mg/kg (∼18 mg/kg ATO) IM dose in rats and >1 month with 11 mg/kg (∼10 mg/kg ATO) IM dose in cyno. No dose dumping with plasma levels higher than C_min_ for >70 days showcased the superiority of mCKX352 to mCBE161 for the same equivalent dose of 10 mg/kg ATO. The duration of chemoprotection in both oiland aqueousbased systems is >1 month in monkeys with a very low dose of 10 mg/kg. This was feasible due to slow release of the ATO into the circulation and no sudden unoptimized high burst. Thus, the ratio of peak (C_max_) and trough (C_min_) concentrations is minimized due to the physical characteristics and optimal formulations of the prodrugs.

While prodrugs enhance the physiochemical properties to enable the desired profile needed for optimal formulation and thus required plasma levels for extended protection, the residual circulating prodrugs could cause adverse events. The presence of prodrugs may also improve half-life of the parent drug and tissue distribution that could impact safety. IM injections with mCKX352 did show evidence of reversible inflammation and muscular necrosis; however, these findings with mCBE161 were not reversible. Importantly, other drugs such as haloperidol decanoate^58^ and paliperidone palmitate^59^ are given via IM injection and both are tolerable. This is an important aspect since ATO has not been dosed parenterally in humans and would be further delineated in longer-term animal toxicology and human safety studies. Given the current findings of the 14-day rat toxicity study and the historical data of ATO, mCKX352 and mCBE161 are anticipated to be well tolerated in humans. However, after comparing PK and safety data generated here, mCBE161 was selected to advance to first-in-human clinical trials with an improved clinical formulation leading to longer duration and reversible findings from tox studies, details of which will be disclosed in future publications.

## CONCLUSIONS

We have developed a long-acting injectable prophylactic for malaria by formulating a prodrug of ATO. Preclinical data show that plasma exposure is maintained above minimal efficacious dose for >70 and >30 days in rats and monkeys, respectively. The release kinetics of prodrug ATO formulations also have reduced initial concentration compared to ATO. These data indicate that our prodrug formulations promote optimal release kinetics with optimized burst effects for long durations, which highlight their potential as first-in-class prophylaxis for malaria. Based on these data, we are pursuing first-inhuman studies to advance our prodrug formulation of ATO, mCBE161, to the clinic.

## EXPERIMENTAL SECTION

### Synthesis of Atovaquone (ATO) Prodrugs

#### Synthesis of ATO prodrugs was substrate dependent. Three methods followed are given below

##### Method A^54^

mCKX352: To a 1,000 mL round bottom flask protected from light, ATO (10 g, 27.5 mol., 1.0 equiv.) and methylene chloride (500 mL) were added. NaOH (330 ml of 10% NaOH solution, 825 mmol., 30 equiv.) was added in one portion and the solution was stirred for 4 hours at room temperature. Due to the formation of a slurry, we used a mechanical stirrer. This was followed by addition of heptanoyl chloride (11 ml, 71.5 mol., 2.6 equiv.) dropwise at room temperature. The reaction mixture was stirred for 24 hours at room temperature. After 24 hours, the organic layer was separated and washed with water (3 × 150 mL) and brine (1 × 150 mL), and the organic phase was dried over MgSO4 and evaporated under reduced pressure. The crude product afforded mCKX352 at 70% yield (9.2 g) after multiple triturations with ethanol. ^1^H NMR (400 MHz, CDCl3) δ 8.15 –8.10 (m, 1H), 8.10 – 8.05 (m, 1H), 7.79 – 7.68 (m, 2H), 7.30– 7.26 (m, 2H), 7.19 – 7.14 (m, 2H), 3.13 – 3.01 (m, 1H), 2.72(t, J = 7.5 Hz, 2H), 2.65 – 2.53 (m, 1H), 2.08 – 1.93 (m, 4H),1.90 – 1.77 (m, 4H), 1.60 – 1.46 (m, 4H), 1.38 (h, J = 3.5 Hz,4H), 0.98 – 0.87 (m, 3H). ^13^C NMR (125 MHz, CDCl3) δ184.55, 178.33, 171.06, 151.52, 145.52, 141.79, 134.10,133.68, 132.40, 131.66, 130.72, 128.48, 128.11, 126.84,126.45, 43.29, 35.95, 34.24, 34.03, 31.48, 29.92, 28.80,24.81, 22.51, 14.03. Anal for C29H31ClO4: Calculated C,72.72; H, 6.52; N, 0. Found: C, 72.83; H, 6.62; N, 0. HRMS (ESI-TOF) [(M-H)–] m/z for C29H31ClO4: Calculated 477.1838; Found: 477.1837

##### Method B^54^

mCBE161: To a 1,000 mL round bottom flask protected from light, ATO (15 g, 41 mol., 1.0 equiv.) and methylene chloride (585 mL) were added. The flask was cooled to 0°C. To this solution was added TEA (8.7 mL, 62.3 mmol, 1.5 equiv.) followed by the drop-wise addition of AcCl (4.4 mL, 62 mmol, 1.5 equiv.) at 0°C. The reaction mixture was then warmed to room temperature and stirred for 24 hours. After 24 hours, the organic layer was washed with water (3 × 150 mL) and brine (1 × 150 mL), and the organic phase was dried over MgSO4 and evaporated under reduced pressure. The crude product afforded mCBE161 at 85% yield (14.2 g) after multiple triturations with ethanol. ^1^H NMR (400 MHz, CDCl3) δ 8.16 – 8.10 (m, 1H), 8.10 – 8.06 (m, 1H), 7.79 – 7.69 (m, 2H), 7.31 – 7.26 (m, 2H), 7.21 – 7.14 (m, 2H), 3.08 (tt, J = 12.3, 3.5 Hz, 1H), 2.60 (tt, J = 12.3, 3.3 Hz, 1H), 2.46 (s, 3H), 2.07 – 1.93 (m, 4H), 1.88 – 1.78 (m, 2H), 1.61 – 1.48 (m, 4H). ^13^C NMR (125 MHz, CDCl3) δ 184.50, 178.28, 168.14, 151.38, 145.51, 141.83, 134.14, 133.71,132.38, 131.65, 130.66, 128.48, 128.12, 126.86, 126.45,77.25, 77.00, 76.75, 43.27, 35.96, 34.21, 29.94, 20.60. Anal for C24H21ClO4: Calculated C, 70.50; H, 5.18; N, 0. Found: C, 70.23; H, 4.94; N, 0. HRMS (ESI-TOF) [(M-H)–] m/z for C24H21ClO4: Calculated 407.1056; Found: 407.1054

##### Method C

DHACOCl: To a 500 mL round bottom flask under an inert atmosphere and protected from light DHA (14.5 g, 44 mmol, 1.0 equiv.) DMF (∼5 drops) and anhydrous methylene chloride (315 mL) were added. The reaction was cooled to 0°C and oxalyl chloride (4.5 mL, 53 mmol, 1.2 equiv.) was added dropwise. The solution was stirred at 0°C for 3.5 hours. Upon completion, the solvent was removed in vacuo and used in the next step without further purification.

mCBK068: To a 500 mL round bottom flask protected from light, ATO (8.0 g, 22 mmol, 1.0 equiv.) and wet methylene chloride (300 mL: prepared by shaking 1:1 methylene chloride and water in a separatory funnel and draining off the methylene chloride) were added. Potassium carbonate (7.6 g, 35 mmol., 2.5 equiv.) was added in one portion followed by DHACOCl (15.27 g, 44 mmol, 2.0 equiv.) dropwise as a solution in methylene chloride (80 mL). The solution was stirred at room temperature for 5 hours. Upon completion, the solids were removed via filtration through a pad of Celite. The filtrate was concentrated in vacuo and purified by silica gel chromatography (using a 5% to 25% ethyl acetate in hexanes as eluent) which afforded pure mCBK068 as a pale-yellow oil in 93% isolated yield (8.5 g). ^1^H NMR (400 MHz, CDCl3) δ 8.12 (dd, J = 6.9, 1.9 Hz, 1H), 8.07 (dd, J = 6.8,2.0 Hz, 1H), 7.74 (qt, J = 7.6, 3.8 Hz, 2H), 7.28 (d, J = 2.0 Hz,2H), 7.19 – 7.13 (m, 2H), 5.56 – 5.47 (m, 2H), 5.45 – 5.25 (m,10H), 3.13 – 3.01 (m, 1H), 2.97 – 2.71 (m, 12H), 2.65 – 2.54 (m, 3H), 2.03 (dt, J = 33.6, 9.3 Hz, 6H), 1.82 (d, J = 12.7 Hz,2H), 1.51 (d, J = 13.2 Hz, 2H), 0.96 (t, J = 7.5 Hz, 3H). ^13^C NMR (126 MHz, CDCl^3^) δ 184.50, 178.23, 170.44, 151.41, 145.48,141.83, 134.14, 133.71, 132.35, 132.01, 131.63, 130.64,129.96, 128.55, 128.47, 128.46, 128.28, 128.10, 128.03,127.86, 127.84, 127.37, 127.00, 126.86, 126.45, 43.24,35.87, 34.19, 33.91, 29.89, 25.69, 25.64, 25.62, 25.60, 25.52,22.61, 20.54, 14.25. Anal for C44H49ClO4: Calculated C,78.03; H, 7.29; N, 0. Found: C, 77.82; H, 7.26; N, 0. HRMS(ESI-TOF) [(MH)+] m/z for C44H49ClO4: Calculated 677.3392; Found: 677.3390

### Aqueous (pH) Solubility Measurement

Approximately ∼25 mg of each test material was weighed into a 3 mL vial, followed by the addition of 2.5 mL of each test media (different pH solutions) into the vials. The vials were then capped and stirred at 400 rpm at 25 ºC. The vials were sampled at 24 hours, 48 hours, and 7 days. At each time point, about 0.8 mL suspension was extracted and centrifuged (10000 rpm, room temperature, 4 min). The supernatant liquid was filtered (0.45 μm PTFE filter) and tested by HPLC for solubility (the first few drops of filtrate were used for pH test). The solids were analyzed by X-ray powder diffraction (XRPD).

### Oil Solubility Measurement

Approximately 100∼200 mg of starting material was weighed into each 3 mL glass vial, then 1 mL of oil was added into the vials to form suspensions. Thereafter, an additional 1 mL of each the oils was added if the solids were not completely dissolved. The vials were capped and then stirred at 400 rpm at 25 ºC for the next 48 hours. The vials were sampled at 48 hours and 7 days. At each time point, about 0.8 mL of the suspension was extracted and centrifuged (10000 rpm, room temperature, 4 min). The supernatant liquid was filtered (0.45 μm PTFE filter) and tested by HPLC for solubility (the first few drops of filtrate are discarded). A dilution by THF is done before solubility test. The isolated solids were analyzed by XRPD.

### Oil Solution and Aqueous Suspension Generation

#### Solution

In order to prepare 100 mg/ml solution of mCKX352, 100 mg of the prodrug was weighed into a 5 mL glass vial, then 1.0 mL of sesame oil was added into the vial to dissolve the solids completely. To affect dissolution, four cycles of sonication (1 min), and vortexing (1 min) were carried out followed by heating at 37ºC for 2 min. The vials were either stored at room temperature or at 37ºC. In the case of mCBK068, 100 mg/ml of the solution was made by dissolving 100 mg of prodrug into the 917 μL of the sesame oil given that the density of the prodrug is 1.2 g/mL. A quick sonication for 2 min was used to affect complete dissolution.

#### Suspension

Approximate 93 mg of mCBE161 was weighed into a vial, and it was suspended in 0.5%CMC-Na/0.2%Tween at targeted drug loading of 200 mg/mL. The sample was then stirred at 700 rpm using a magnetic stirrer at 25 ºC for 24 h. The homogenous suspension was collected and analyzed using HPLC and ZPPS. Similar procedure was used for the generation of the ATO suspension.

### Accelerated Stability Studies of Oil solution and Aqueous Suspension

#### Solution

Approximately 30∼37.5 mg of the starting material was weighed into each 5-mL glass vial, then 2.5 mL of oil was added into the vials to dissolve the solids completely at 25ºC. The vials were capped and then stirred at 400 rpm at 25ºC for next 48 h. The vials were sampled at 48 hours and initial purity was determined. Thereafter, approximately 0.8 mL of each solution was pipetted into each vial, and then the vials were placed under the three conditions (25ºC/60%RH, 40ºC/75%RH and 40 ºC/ambient). The vials were then sampled at 7 days and 4 weeks. A dilution by THF is done before purity test.

#### Suspension

Approximate 93 mg of mCBE161 was weighed into a vial, and it was suspended in 0.5%CMC-Na/0.2%Tween at targeted drug loading of 200 mg/mL. The sample was then stirred at 700 rpm using a magnetic stirrer at 25ºC for 24 h. The homogenous suspension was collected and analyzed using HPLC and ZPPS at initial. Then the suspension was stored at RT, 40ºC and 60ºC for accelerated stability test.

### pH Stability Studies

Approximately 0.5∼1 mg of test materials was weighed into a 10-mL volumetric flask followed by the addition of each test media (different pH solutions) into the vials to dissolve the solids completely and then was diluted to the volume. The vials were then capped and kept in stability chambers at 25ºC, and were sampled at 2 h, 1 day, and 7 days to test purity at each time point.

### Bioanalytical Methods of ATO and its Prodrugs

#### kAAC207 (ATO)

A bioanalytical method has been developed for kAAC207 utilizing LC/MS/MS. The following is the detailed procedure. Blood samples from male SD rats were collected in tubes containing K_2_EDTA and centrifuged to obtain plasma. 5 μL aliquots of plasma were treated with 100 μL of ice-cold acetonitrile containing internal standards (100 ng/mL Labetalol & 100 ng/mL dexamethasone & 100 ng/mL tolbutamide & 100 ng/mL verapamil & 200 ng/mL diclofenac & 100 ng/mL celecoxib) for liquid-liquid extraction of analyte, kAAC207. Samples were vortexed then centrifuged at 13000 rpm for 5 min. Samples were kept at 4°C during centrifugation. 2 μL (injection volume) of supernatant was injected into a Waters Acquity UPLC system fitted with a Waters UPLC Protein BEH C4 50 × 2.1 mm, 1.7 μm column maintained at 50°C. Mobile phase A consisted of 0.1% formic acid and 2 mM ammonium formate in 95% water/5% acetonitrile (v/v). Mobile phase B consisted of 0.1% formic acid and 2 mM ammonium formate in 5% water/95% acetonitrile (v/v). Flow rate was maintained at 0.6 mL/min. A gradient up to 1.5 min was used for elution of the analyte and IS. Starting with 80% of mobile phase A, mobile phase B increased to 95% at 0.6 min. Mobile phase B was held at 95% until 1.41 min, where the gradient reverted to 80% mobile phase A. Concentrations were quantitated in plasma samples using an AB SCIEX 6500+ using Turbo Spray in negative mode. Detection was performed by single reaction monitoring (kAAC207, 365.1 → 337.3; (IS) Diclofenac, 294.2 → 250). A calibration curve to quantify blood samples was generated for kAAC207 in blank male SD rat plasma consisting of a linear range from 2-500 ng/mL. The analytical lower limit of quantitation (LLOQ) for kAAC207 in plasma was typically ∼ 1-2 ng/mL and accuracy, precision and recovery were within acceptable limits.

#### mCBE161

A bioanalytical method has been developed for mCBE161 utilizing LC/MS/MS. The following is the detailed procedure. Blood samples from male SD rats were collected in tubes containing K_2_EDTA and centrifuged to obtain plasma. 50 μL aliquots of plasma were treated with 200 μL of ice-cold acetonitrile containing internal standards (100 ng/mL Labetalol & 100 ng/mL dexamethasone & 100 ng/mL tolbutamide & 100 ng/mL verapamil & 100 ng/mL glyburide & 100 ng/mL celecoxib) for liquid-liquid extraction of analyte, mCBE161. Samples were vortexed then centrifuged at 13000 rpm for 5 min. Samples were kept at 4°C during centrifugation. 10 μL (injection volume) of supernatant was injected into a Waters Acquity UPLC system fitted with a Waters UPLC Protein BEH C4 50 × 2.1 mm, 1.7 μm column maintained at 50°C. Mobile phase A consisted of 0.1% formic acid and 2 mM ammonium formate in 95% water/5% acetonitrile (v/v). Mobile phase B consisted of 0.1% formic acid and 2 mM ammonium formate in 5% water/95% acetonitrile (v/v). Flow rate was maintained at 0.5 mL/min. A gradient up to 4.0 min was used for elution of the analyte and IS. Starting with 90% of mobile phase A, mobile phase B increased to 95% at 3.5 min. Mobile phase B was held at 95% until 3.81 min, where the gradient reverted to 90% mobile phase A. Concentrations were quantitated in plasma samples using an AB SCIEX 6500+ using Turbo Spray in negative mode. Detection was performed by single reaction monitoring (mCBE161, 408.2 → 367.2; (IS) Tolbutamide, 269.0 → 169.8). A calibration curve to quantify blood samples was generated for mCBE161 in blank male SD rat plasma consisting of a linear range from 50-1000 ng/mL. The analytical lower limit of quantitation (LLOQ) for mCBE161 in plasma was typically ∼ 50 ng/mL and accuracy, precision and recovery were within acceptable limits.

#### mCKX352

A bioanalytical method has been developed for mCKX352 utilizing LC/MS/MS. The following is the detailed procedure. Blood samples from male Sprague Dawley rats were collected in tubes containing K_2_EDTA and centrifuged to obtain plasma. 8 μL aliquots of plasma were treated with 80 μL of ice-cold acetonitrile containing internal standards (100 ng/mL labetalol, 100 ng/mL dexamethasone, 100 ng/mL tolbutamide, 100 ng/mL verapamil, 200 ng/mL diclofenac, & 100 ng/mL celecoxib) for liquid-liquid extraction of analyte, mCKX352. Samples were vortexed then centrifuged at 13000 rpm for 5 min. Samples were kept at 4°C during centrifugation. 6 μL (injection volume) of supernatant was injected into a Waters Acquity UPLC system fitted with a Waters UPLC Protein BEH C4 50 × 2.1 mm, 1.7 μm column maintained at 50°C. Mobile phase A consisted of 0.3% formic acid in water. Mobile phase B consisted of 0.3% formic acid in acetonitrile. Flow rate was maintained at 0.6 mL/min. A gradient up to 1.8 min was used for elution of the analyte and internal standard. Starting with 80% of mobile phase A, mobile phase B increased to 80% at 1.2 min. Mobile phase B was held at 80% until 1.71 min, where the gradient reverted to 80% mobile phase A. Concentrations were quantitated in plasma samples using an AB SCIEX 6500+ using Turbo Spray in negative mode. Detection was performed by single reaction monitoring (mCKX352, 478.2 → 366.1; (internal standard) diclofenac, 294.2 → 250.1). A calibration curve to quantify blood samples was generated for mCKX352 in blank male Sprague Dawley rat plasma consisting of a linear range from 5–500 ng/mL. The analytical lower limit of quantitation (LLOQ) for mCKX352 in plasma was typically ∼ 5 ng/mL and accuracy, precision and recovery were within acceptable limits.

### Drug Administration

A summary of ATO and ATO prodrug drug administration parameters is given below (**Tables S1 & S2**). The solutions and suspensions were administered IM at the hind leg of rat, the right/left muscular flank (biceps femoris) of the hind leg of dog, and the left/right muscular flank (biceps femoris) of monkey.

### PK Studies of ATO and its Prodrugs

#### Rats

Prodrugs were dosed IM to male Sprague Dawley rats with three rats per dosing group. All rats were fasted for at least 12 hours prior to the administration. After dosing, 75 μL of blood was collected at predetermined time points (0.021, 0.042, 0.167, 0.333, 1, 2, 3, 4, 5, 6, 7, 14, 21, 28, 42days). Blood samples were processed for plasma by centrifugation at approximately 4°C, 10,000 rpm for 3 min within half an hour of collection. Plasma samples were stored in polypropylene tubes, quick-frozen over dry ice and kept at −70±10°C until LC/MS/MS analysis. Plasma concentration versus time data was analyzed by non-compartmental approaches using the Phoenix WinNonlin 6.3 software program.

#### Dogs

mCBK068 was dosed IM to male Beagle dogs with three dogs per dosing group. Animals were fed twice daily. Stock dogs were fed approximately 220 grams of Certified Dog Diet daily (Beijing Vital Keao Feed Co., Ltd. Beijing, P. R. China). After dosing, 500 μL of blood was collected at predetermined time points (0.010, 0.042, 0.125, 0.208, 0.333, 1, 2, 3, 4, 5, 6, 13, 20, 27 days). Blood samples were collected into a commercial tube (Jiangsu Kangjian medical supplies co., LTD) containing Potassium (K2) EDTA*2H2O (0.85-1.15 mg) on wet ice and processed for plasma. Samples were centrifuged (3,000 x g for 10 minutes at 2 to 8°C) within one hour of collection. The plasma samples about 0.1 mL* 2 aliquots (once for BA, and the other for back up) were transferred into labeled polypropylene micro-centrifuge tubes and stored frozen in a freezer set to maintain −60°C or lower until bio analysis.

#### Monkeys

Prodrugs were dosed IM to male cynomolgus monkeys with three monkeys per treatment group. Animals were fed twice daily. Stock monkeys were fed approximately 120 grams of Certified Monkey Diet daily (Beijing Keao Xieli Feed Co., Ltd. Beijing, P. R. China). After dosing, 500 μL of blood was collected at predetermined time points (0.010, 0.042, 0.125, 0.333, 1, 2, 3, 4, 5, 6, 7, 14, 21, 28, 42, 55, 72, 103, 134 days). Blood samples were collected into a commercial tube (Jiangsu Kangjian medical supplies co., LTD) containing Potassium (K2) EDTA*2H_2_O (0.85-1.15 mg) on wet ice and processed for plasma. Samples were centrifuged (3,000 x g for 10 minutes at 2 to 8°C) within one hour of collection. The plasma samples about 0.1 mL* 2 aliquots (once for BA, and the other for back up) were transferred into labeled polypropylene micro-centrifuge tubes and stored frozen in a freezer set to maintain −60°C or lower until bio analysis.

#### Two-week Rat Toxicity Study

The study was designed to assess the potential systemic toxicity, target organs, and TK profile of mCMX352 or mCBE161, when administered twice (on days 1 and 14) via IM route at dose levels of 20, 100, and 200 mg/kg mCMX352 or 20, 80, and 250 mg/kg mCBE161. For each of the test articles, a total of 41 male and 41 female Sprague Dawley rats were randomly assigned to 4 main groups (G1 to G4; 5 rats/sex/group) and 4 TK groups (vehicle: 3 rats/sex/group and test article treated: 6 rats/sex/group). The test articles were formulated in vehicle (mCBE161: 0.5% CMC and 0.2% tween 80 in water and mCMX352: sesame oil) and dosed via IM injection in the thigh muscle once on days 1 and 14 at a dose volume of 0.667 mL/kg. The vehicle control group received sesame oil vehicles on respective days. The dose formulations were prepared for day 1 and day 14 of the study and were analyzed for mCB161 or mCMX352 content using a validated HPLC method.

Observations included morbidity/mortality check, clinical signs of toxicity, detailed clinical examination, injection site evaluation, body weight, and food consumption. Blood samples at scheduled time points from all TK groups were collected (day-1 and day-14 samples were collected at predose (0 hours), 0.25, 1, 4, 24, 72, 120, 168, and 240 hours post dose) to determine the plasma exposure and TK profiles of mCBE161 and ATO or mCMX352 and ATO. TK data analysis was performed in Phoenix WinNonlin (version 6.3). TK parameters such as Tmax, Cmax, AUC_0-240h_, AUC_inf_, and terminal plasma half-life (t_1/2_) were reported.

At the end of the treatment period (day 15), hematology, clinical chemistry, urinalysis, gross pathology, and organ weight measurements were performed. Histology was performed on all preserved organs for vehicle control and high dose groups and site of injections (thigh muscle and skin with subcutaneous tissue).

## ASSOCIATED CONTENT

### Supporting Information

Scheme outlining ATO prodrug synthesis and Tables summarizing physical properties of prodrugs, rat PK, and cyno PK (PDF):

- Supplementary Figure S1: Synthesis of ATO prodrugs
- Supplementary Table S1: Summary of physical properties of ATO and ATO prodrug lead compounds
- Supplementary Figure S2: Rat pharmacokinetics of ATO and ATO prodrugs administered via IM depot
- Supplementary Figure S3: Cyno PK studies of ATO prodrugs administered via IM depot This material is available free of charge via the Internet at http://pubs.acs.org

## AUTHOR INFORMATION

### Author Contributions

The manuscript was written through contributions of all authors. All authors have given approval to the final version of the manuscript.

## Notes

Any additional relevant notes should be placed here.

## ACKNOWLEDGMENT

We would like to acknowledge Dr. Kit Bonin and Dr. Geneva Hargis for writing and graphic support.

## ABBREVIATIONS

API: Active pharmaceutical ingredient
ATO: Atovaquone
BQL: Below quantitation limit
C_last_: Final concentration measured
clogP: Calculated log partition
C_max_: Maximal concentration
CMC: Carboxymethyl cellulose
C_min_: Minimum efficacious concentration
DADDS: 4-4’-diacetylaminosulphone
DHA: Docosahexaenoic acid
EVA: Ethylene-vinyl acetate
GLP: Good laboratory practice
GRAS: Generally recognized as safe
IM: Intramuscular
IPTi: intermittent preventative treatment in infancy
IPTp: Intermittent preventative treatment in pregnancy
IRS: Indoor residual spraying
ITNs: Insecticide-treated bed nets
LAI: Long-acting injectable
MMV: Medicines for Malaria Venture
NOAEL: No observed adverse event level
PK: Pharmacokinetic
PO: oral
PrEP: Pre-Exposure Prophylaxis
SDG: Sustainable development goals
SLS: Sodium laureth sulfate
SMC: Seasonal malarial chemoprevention
SPAQ: Sulfadoxine-pyrimethamine plus amodiaquine
TCP: Target candidate profile
TI: Therapeutic index
TK: Toxicokinetics
T_max_: Time to peak drug concentration
TPP: Target produce profile
UN: United Nations
WHO: World Health Organization
WRAIR: Walter Reed Army Institute of Research

Authors are required to submit a graphic entry for the Table of Contents (TOC) that, in conjunction with the manuscript title, should give the reader a representative idea of one of the following: A key structure, reaction, equation, concept, or theorem, etc., that is discussed in the manuscript. Consult the journal’s Instructions for Authors for TOC graphic specifications.

## Insert Table of Contents artwork here

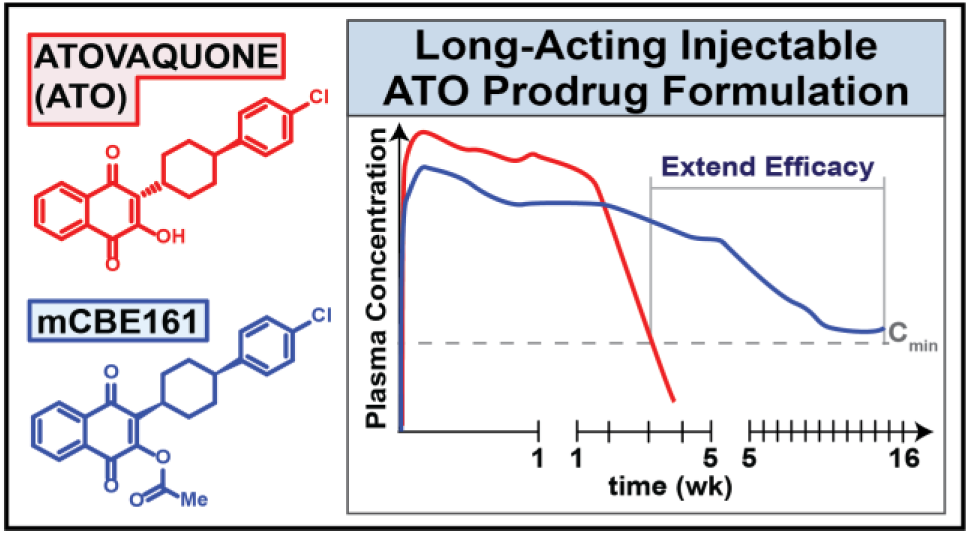

