## Supporting Info for "A Prodrug Strategy to Reposition Atovaquone as a Long-Acting Injectable for Malaria Chemoprotection" (from another version of this preprint: version 1, 2024-02-12)

Table of Figures:

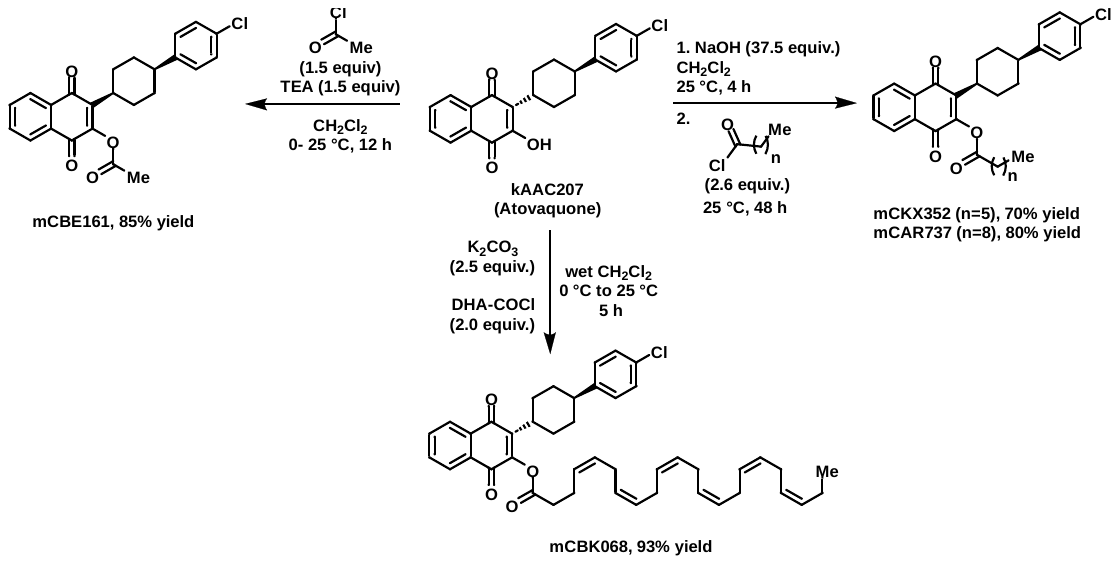

**Figure S1. Synthesis of ATO Prodrugs**

**Table S1. Summary of physical properties of ATO and ATO prodrug lead compounds.**

| **Physiochemical Properties** | | **kAAC207 (ATO)** ^a^ | **mCBE161** ^a^ | **mCKX352** ^a^ | **mCBK068** ^a^ |
| --- | --- | --- | --- | --- | --- |
| Molecular Weight | | 366.84 | 408.88 | 479.01 | 677.31 |
| Melting point by DSC (°C) | | 222.8 | 163.2 | 76.1 | NA ^b^ |
| Solubility at pH 2 (HCl in water) (µg/mL) | 24 h | <3.8 | <4.5 [<4.0] | <1.08 [0.83] | <11.4 [<6.2] |
|  | 48 h | <3.8 | <4.5 [<4.0] | <1.08 [0.83] | <11.4 [<6.2] |
|  | 168 h | <3.8 | <4.5 [<4.0] | <1.08 [0.83] | <11.4 [<6.2] |
| Solubility at pH 4 (AcOH and NaOH in water) (µg/mL) | 24 h | <3.8 | <4.5 [<4.0] | <1.08 [0.83] | <11.4 [<6.2] |
|  | 48 h | <3.8 | <4.5 [<4.0] | <1.08 [0.83] | <11.4 [<6.2] |
|  | 168 h | <3.8 | <4.5 [<4.0] | <1.08 [0.83] | <11.4 [<6.2] |
| Solubility at pH 6 (AcOH and NaOH in water) (µg/mL) | 24 h | <3.8 | <4.5 [<4.0] | <1.08 [0.83] | <11.4 [<6.2] |
|  | 48 h | <3.8 | <4.5 [<4.0] | <1.08 [0.83] | <11.4 [<6.2] |
|  | 168 h | <3.8 | <4.5 [<4.0] | <1.08 [0.83] | 0.12 [0.06] |
| Solubility at pH 8 (Tris and HCl in water) (µg/mL) | 24 h | <3.8 | <4.5 [<4.0] | <1.08 [0.83] | <11.4 [<6.2] |
|  | 48 h | <3.8 | <4.5 [<4.0] | <1.08 [0.83] | <11.4 [<6.2] |
|  | 168 h | <3.8 | <4.5 [<4.0] | <1.08 [0.83] | 0.08 [0.04] |
| Solubility in fractionated coconut oil at 25 °C (mg/mL) | 48 h | ND | ND | 68.6 [52.6] | >135 [>85] |
|  | 168 h | ND | ND | 77.6 [59.4] | ND |
| Solubility in soybean oil at 25 °C (mg/mL) | 48 h | ND | ND | 41.6 [31.8] | ND |
|  | 168 h | ND | ND | 46.9 [35.9] | ND |
| Solubility in olive oil at 25 °C (mg/mL) | 48 h | ND | ND | 43.0 [32.9] | ND |
|  | 168 h | ND | ND | 46.8 [35.8] | ND |
| Solubility in castor oil at 25 °C (mg/mL) | 48 h | ND | ND | 30.7 [23.5] | ND |
|  | 168 h | ND | ND | 37.8 [28.9] | ND |
| Solubility in sesame oil at 25 °C (mg/mL) | 48 h | 1.3 | 4.3 [3.9] | ND | 34.9 [18.9] |
|  | 168 h | 1.2 | 3.9 [3.5] | ND | 32.9 [17.8] |
| Solubility in Miglyol 812 at 25 °C (mg/mL) | 48 h | 3.1 | 6.4 [5.7] | ND | >283.6 [>153.6] |
|  | 168 h | 2.8 | 7.7 [6.9] | ND | 212.5 [115.1] |
| Maximum drug Loading (mg/mL) | Conc/ loading  (mg/mL) | 203 | 200 [179] | 200 [153] | 1200 [650] |
|  | Vehicle | 1% (w/v) F108, 0.2% (w/v) SLS | 0.5% (w/v) CMC, 0.2% (w/v) Tween 80 | 20% (v/v) benzyl benzoate, 80% (v/v) fractionated coconut oil | Sesame oil |
|  | Type of formulated material | Suspension | Suspension | Solution | Solution |

*^a^ ND = not determined; Data in brackets are the values normalized to ATO concentration/loading in solution/suspension;;*

*^b^ mCBK068 is an oil at room temperature.*

**Table S2.** **Rat pharmacokinetics of ATO and ATO prodrugs administered via IM depot**.

| **Rats^a^** | **IM ATO** | **IM mCBE161** | **IM mCKX352** | **IM mCBK068** |
| --- | --- | --- | --- | --- |
| Dose of Prodrug (mg/kg) | NA | 20 | 20 | 40 |
| Loading of Prodrug (mg/ml) | NA | 240 | 227 | 545 |
| Dose of ATO (mg/kg) | 20 | 18 | 15 | 21 |
| Loading of ATO (mg/ml) | 200 | 215 | 174 | 295 |
| Dose volume (ml/kg) | 0.1 (25 µL) ^b^ | 0.08 (21 µL) ^b^ | 0.09 (26 µL) ^b^ | 0.07 (18 µL) ^b^ |
| Formulation | 1% (w/v) F108 and 0.2% (w/v) SLS | 0.1% (w/v) F108 and 0.2% (w/v) SLS | Sesame oil | Sesame oil |
| Duration of study (wks) | 4 | 16 | 16 | 13 |
| C_last_ (nM) | 19 (ND) | 135 (ND) | 122(64) | 2,256 (935) |
| C_max_ (nM) | 35,532 (5682) | 11,149 (2007) | 2,058 (335) | 13,812 (4909) |
| C_max_/Dose (nM/mg/kg) | 1,777 (284) | 619 (112) | 137 (22) | 658 (234) |
| C_max_ /C_min_ (nM)^c^ | 130.6 | 40.9 | 7.6 | 50.8 |
| AUC_0-inf_ (nM*h) | 5,569,351 (489,767) | 3,646,565 (213,498) | 1,558,238 (323,606) | 15,834,303 (3,226,253) |
| AUC_0-inf_ /Dose (nM*h/mg/kg) | 278,468 (24,488) | 202,587 (11,861) | 103,883 (21,574) | 754,014 (153,361) |
| AUC_0-last_ (nM*h) | 5,569,959 (510,346) | 3,586,680 (307,037) | 1,436,850 (268,220) | 13,335,405 (1,900,598) |
| AUC_0-last_ /Dose (nM*h/mg/kg) | 278,498 (25,517) | 199,260 (17,058) | 95,790 (17,881) | 635,019 (90,505) |
| T_1/2_ (d) | 2.2 (0.4) | 12.3 (6) | 28 (1) | 30.5 (5.5) |
| T_max_ (d) | 1.3 (0.6) | 1 (0) | 6.7 (0.6) | 10.7 (15) |
| MRT_0-last_ (d) | 5.6 (0.7) | 18.9 (3.4) | 28 (2) | 33.8 (4.3) |
| CL/F (mL/min/kg) | 0.16 (0.02) | 0.22 (0.01) | 0.46 (0.1) | 0.12 (0.02) |
| LLOQ of ATO (nM) | 2.7 (first 3 wks)  5.5 (last wk) | 5.5 (first 2 wks)  2.7 (last 14 wks) | 5.5 | ND |
| LLOQ of Prodrug (nM) | Not Applicable | ND | 10.4 | ND |

*^a^ AUC, area under the plasma concentration–time curve; AUC_0-inf_, AUC to infinity; AUC_0-last_, AUC to time of last measurable concentration, C_last_, last quantifiable plasma concentration at last; C_max_, maximum plasma concentration; SD, standard deviation is given in parentheses; T1/2, terminal elimination half‐life; NA = not applicable; ND = not determined; The doses of equivalent ATO has been used to calculate dose normalized parameters.*

*^b^ Dose volume in parenthesis is calculated assuming the mean mass of a rat to be 0.25 kg.*

*^c^ C_min_ = 272 nM*

**Table S3. Cyno PK studies of ATO prodrugs administered via IM depot.**

| **Cynos^a^** | **IM mCBE161** | **IM mCKX352** | **IM mCBK068** |
| --- | --- | --- | --- |
| Dose of Prodrug (mg/kg) | 11 | 13 | 19 |
| Loading of Prodrug (mg/ml) | 100 | 100 | 190 |
| Dose of ATO (mg/kg) | 10 | 10 | 10 |
| Loading of ATO (mg/ml) | 90 | 77 | 103 |
| Dose volume (ml/kg) | 0.11 (292 µL) ^b^ | 0.13 (425 µL) ^c^ | 0.1 (300 µL) ^d^ |
| Formulation | 0.5% (w/v) CMC-Na /0.2% (w/v) Tween 80 | Sesame oil | Sesame oil |
| Duration of study (wks) | 8 | 24 | 19 |
| C_last_ (nM) | 6 (ND) | 54 (40) | 77 (47) |
| C_max_ (nM) | 2,890 (295) | 937 (392) | 1,071 (236) |
| C_max_/Dose (nM/mg/kg) | 289 (30) | 94 (39) | 107 (24) |
| C_max_ /C_min_ (nM)^e^ | 10.7 | 3.5 | 4.0 |
| AUC_0-inf_ (nM*h) | 1,091,972 (153,679) | 1,339,361 (468,806) | 957,704 (127,658) |
| AUC_0-inf_ /Dose (nM*h/mg/kg) | 109,197 (15,368) | 133,936 (46,881) | 95,770 (12,766) |
| AUC_0-last_ (nM*h) | 1,091,277 (153,630) | 1,241,440 (456,018) | 832,060 (27,422) |
| AUC_0-last_ /Dose (nM*h/mg/kg) | 109,128 (15,363) | 124,144 (45,602) | 83,206 (2,742) |
| T_1/2_ (d) | 3.3 (0.4) | 48 (9) | 44 (8) |
| T_max_ (d) | 14 (0) | 16 (11) | 5 (7) |
| MRT_0-last_ (d) | 16.7 (1.7) | 48 (9) | 42 (10) |
| CL/F (mL/min/kg) | 0.42 (0.06) | 0.37 (0.16) | 0.49 (0.06) |
| LLOQ of ATO (nM) | 2.7 | 5.5 | 8 (first 10 wks)  5.5 (for 14^th^ and 19^th^ wk) |
| LLOQ of Prodrug (nM) | 122 | 10 | 74 |

*^a^ AUC, area under the plasma concentration–time curve; AUC_0-inf_, AUC to infinity; AUC_0-last_, AUC to time of last measurable concentration, C_last_, last quantifiable plasma concentration at last; C_max_, maximum plasma concentration; SD, standard deviation is given in parentheses; T_1/2_, terminal elimination half‐life; ND = Not Determined*

*^b^ Dose volume in parenthesis is calculated based on the average weight of a monkey to be 2.65 kg.*

*^c^ Dose volume in parenthesis is calculated based on the average weight of a monkey to be 3.27 kg.*

*^d^ Dose volume in parenthesis is calculated based on the average weight of a monkey to be 3.00 kg.*

*^e^ C_min_ = 272 nM*
